# Hydrodynamic and electrophoretic properties of Trastuzumab/HER2 complexes. Experiments and modeling

**DOI:** 10.1101/483081

**Authors:** J. Ramos, J.F. Vega, V. Cruz, E. Sánchez-Sánchez, J. Cortés, J. Martínez-Salazar

## Abstract

The combination of hydrodynamic and electrophoretic experiments and computer simulations is indeed a powerful approach to study the interaction between proteins. In this work, we present hydrodynamic and electrophoretic experiments in aqueous solution along with molecular dynamics and hydrodynamic modeling to monitor and compute biophysical properties of the interactions between the extracellular domain of the HER2 protein (eHER2) and the monoclonal antibody trastuzumab (TZM). The importance of this system relies on the fact that the overexpression of HER2 protein is related with the poor prognosis breast cancers (HER2++ positives) being the TZM a monoclonal antibody for the treatment of this cancer. We have found and characterize two different complexes between the TZM and eHER2 proteins (1:1 and 1:2 TZM:eHER2 complexes). The conformational features of these complexes regulate their hydrodynamic and electrostatic properties. Thus, the results indicate a high degree of molecular flexibility in the systems, that ultimately leads to higher values of the intrinsic viscosity and as well as lower values of diffusion coefficient than those expected for simple globular proteins. A highly asymmetric charge distribution is detected for the monovalent complex (1:1 complex), which has strong implications in correlations between the experimental electrophoretic mobility and the modeled net charge. In order to understand the dynamics of these systems and the role of the specific domains involved, it is essential to find biophysical correlations between dynamics, macroscopic transport and electrostatic properties. The results should be of general interest for researchers working in this area.

## 1. Introduction

HER2 (or ErbB2), one of the epidermal growth factor receptors family (EGFR), is well-known to be overexpressed in aggressive human breast cancer. Remarkably, HER2 is the unique EGFR which is activated without ligand. Thus, high amount of HER2 on the cell surface promotes the dimer formation with HER2 (homodimers) or other member of the EFGRs family (heterodimers). These dimers lead to the phosphorylation of the intracellular tyrosine kinase domains, which trigger several signaling pathways related to cellular oncogenic processes (i.e. cellular proliferation, survival, motility, or angiogenesis)(1). For these reasons, e-HER2 is an attractive strategy as a therapeutic protein target. In the late 1990s, trastuzumab (TZM), a monoclonal antibody, showed significant anti-HER2 efficiency in the clinical treatment of tumors, in combination with chemotherapy agents(1–4). TZM binds specifically to an epitope located on e-HER2 domain IV(5, 6). The mechanism of action of TZM is not yet well understood(7). Several mechanisms of action of TZM have been proposed: i) induction of antibody-dependent cellular cytotoxicity, ii) prevention of e-HER2 domain cleavage and finally the most important in HER-2 overexpressed cancers, iii) the dimerization inhibition which prevents the cell growth and proliferation(8). Irrespective of the precise mechanism involved, the overall result is two-fold: an increase of the apoptosis and a suppression of the cell proliferation.

Several crystal structures of antigen-binding Fab domains have been already elucidated-trastuzumab:e-HER2(9), pertuzumab:e-HER2(10), cetuximab:e-EGFR(11), matuzumab:e-EGFR(12) complexes with PDB IDs: 1N8Z, 1S78, 1YY8 and 3C09, respectively-, to cite only some examples from the RCSB-PDB database(13). All this work has contributed to improve the understanding of the binding region between antibody and receptor.

Important features of proteins and biomacromolecular complexes are their sizes and shapes. Both can be obtained through hydrodynamic properties. Those properties, have been investigated through a combination of experimental techniques such as sedimentation, gel filtration, viscosimetry, light scattering as well as electron microscopy. Interesting reviews about the results were published by Harding (14, 15). It is well known that the hydrodynamics of proteins may be altered by ligands (16), phospholipids(17), polyelectrolites(18) and by the formation of both complexes(19),(20) and assemblies(21). The importance of protein hydrodynamics is also especially interesting in the field of biotechnology. As an example we may cite the study of protein PEGylation (22), and the protein or antibody formulation and aggregation (23–25). Experiments in solution of the formation of complexes between different extracellular domains of the EGFR family has been reported by Ferguson et al.(26)

In the context of the above discussion the combination of computer simulations and experiments are proved to be a valuable approach to sample various aspects of the conformational properties of proteins and possible interactions between them(14). For instance, Garcia de la Torre et al. have revised the ability of hydrodynamic modelling to study the protein conformations in solution, using two kinds of “coarse-grained” approximations: the whole-body ellipsoid-based modelling (for globular proteins) and the bead-modelling models (suitable for more complex structures)(27). They have also implemented these models in a software suite to study complex protein structures (i.e. antibodies(28) or protein complexes(29)) with a high capability of prediction(30). Rai et al. proposed the SOMO (SOlution MOdeller) approach, based on a direct correspondence between the atom positions in a macromolecule and the coarse-grained bead model that is used to calculate the hydrodynamic properties(31). This method is implemented within the Ultra-Scan suite(32). Finally, a rather different approximation was proposed by Aragon et al.(33). The method is based on a precise boundary element numerical solution of the exact formulation of the hydrodynamic resistance problem with stick boundary conditions (BEST). The coarse-grained BEST models are obtained from full atomistic molecular dynamics simulations in a multiscale framework. This approach along with experimental measurements has been successfully applied by Brandt et al. to study the case of the conformational space of a monoclonal IgG antibody in solution (34). Concerning this particular point, we have explored, in a previous paper, the solution properties of e-HER2 monomer and its homodimer, here combining hydrodynamic experiments and computer simulations based on BEST(35). For the homodimer, we also characterized the interactions between the two protomers. In the present work, our study is focused on the characterization of the interaction between the e-HER2 protein and the TZM in solution. The combination of dynamic light scattering (DLS), size exclusion chromatography (SEC) and Z-potential measurements (electrophoretic mobility) methods along with multi-scale computational models has emerged in a detailed description of the important properties of these protein complexes.

## 2. Materials and methods

### 2.1 Experimental details

The TZM Herceptin^©^ (stock solution at 21 mg·ml^−1^) was kindly provided by one of us (JC) from Hospital Ramón y Cajal (Madrid, Spain). On the other hand, Sino Biological, Inc. (Beijing, China) provided the glycosylated extracellular HER2 tagged with a 10-length polyhistidine peptide on the C-terminal (g-eHER2-his). These samples were prepared using human cells, in which a DNA sequence encoding the extracellular domain (Met 1-Thr 652) of human HER2 (NCBI Reference Sequence: NP_004439.2) was used. The theoretical molecular mass was 71kDa, but because a glycosylation, the samples migrate to a weight of 100-110 kDa in SDS-PAGE under reducing conditions. After multi-step purification process based on chromatography and filtration, the purified products can reach at least 95% purity. It is stored as a lyophilised powder from sterile PBS (pH 7.4). Before lyophilisation, a combination of trehalose and mannitol in a concentration of 5 to 8% are added as protectants.

Desalting and buffer exchange have been carried out using centrifugal concentrators (Amicon Ultra-0.5 ml, Merck Millipore, Billerica, USA). Both g-eHER2-his and TZM antibody samples were finally filtered (Millex-GV 0.22 μm, Merck Millipore, Billerica, USA) and stored in 20 mM Tris-HCl pH 7.5, 150 mM NaCl to subsequent analysis of hydrodynamic and electrophoretic properties and preparation of the mixtures for the selected molar ratios, i.e., TZM/HER2 2:1 and 1:2. Water for all buffers and dilution was obtained from Milli-Q water purification system (Merck Millipore, Billerica, USA). In order to verify the correct functioning of the equipment we have used a set of globular proteins (BSA and aldolase, GE Healthcare, Buckinghamshire, UK).

Experimental details concerning size exclusion chromatography (SEC), dynamic light scattering (DLS) and electrophoretic mobility (EM) were reported in our previous articles(21, 35). The sample concentration in both SEC and EM measurements were 1-3 mg·ml^−1^. Meanwhile, for DLS experiments, a concentration of < 1 mg·ml^−1^ was used. In all cases the temperature was 293 K.

### 2.2 Computational simulations

#### 2.2.1 Construction of the models

Two systems that correspond to the interaction between one TZM molecule with either one or two g-eHER2 fragments (1:1 and 1:2 complexes), respectively, have been simulated. Both models were built taking into account the structures of the antibody and the g-eHER2-his reported in our previous work (35). The interaction between the TZM Fabs and the extracellular receptor fragments was assembled using as template the crystallographic structure (PDB code: 1N8Z) (9). The Sybyl-X package (36) was used to build the initial conformations of both systems taking into account the glycosylated fragments and the poly-histidine tag associated to the g-eHER2 C-terminal section. Fig. 1 shows a representation of the initial structures for the 1:1 and 1:2 complexes, respectively. The 1:2 complex was considered because it matches the molecular weight experimentally estimated for the case where g-eHER2-his concentration is in excess.

**Fig. 1.**
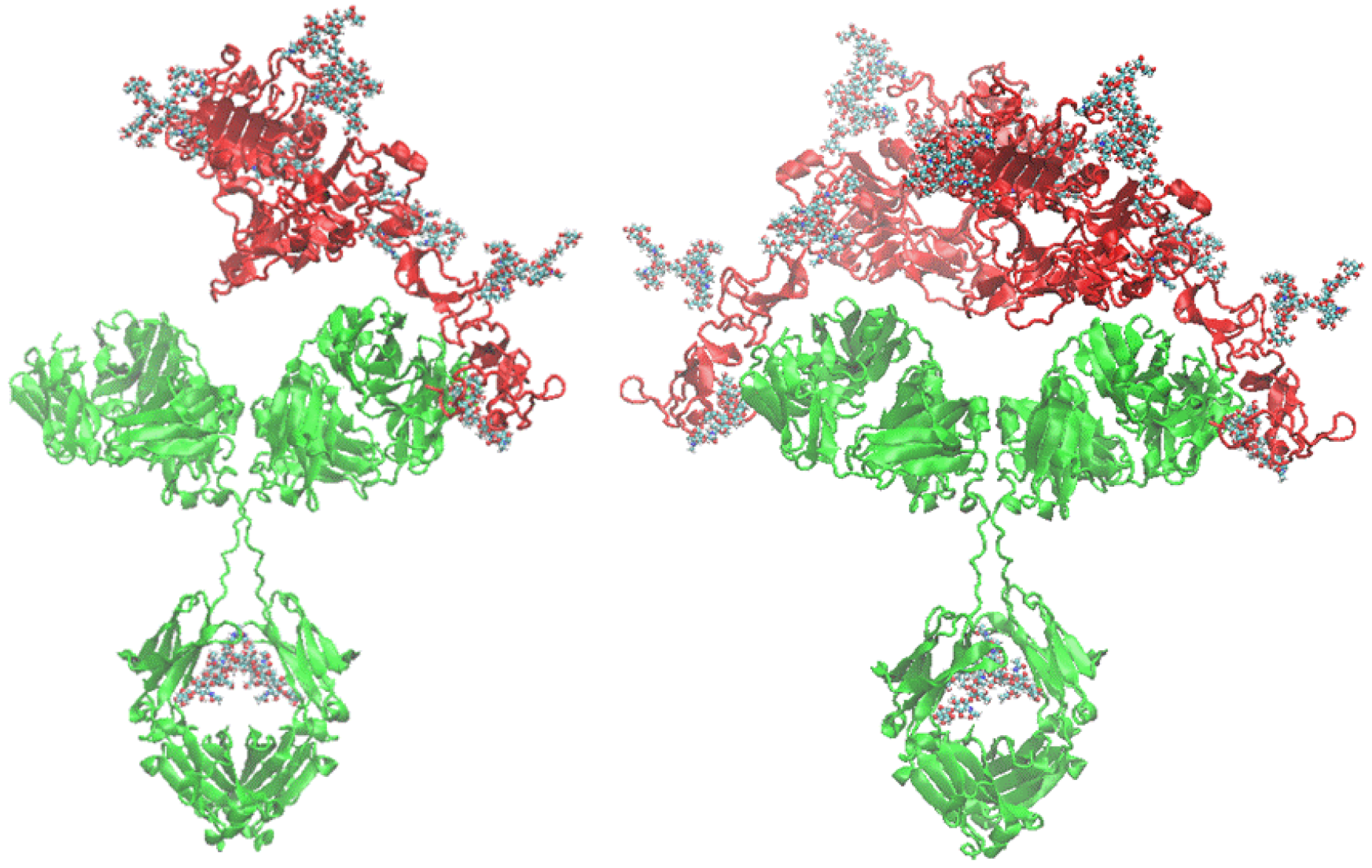
Cartoon representation for the 1:1 (left) and 1:2 (right) complexes. The TZM antibody protein is represented in green and the HER2 ECD in red colors. The glycans are represented as CPK models.

#### 2.2.2 Atomistic molecular dynamics (MD) simulations

MD trajectories for the complexes shown in Fig. 1 were obtained using the Amber 14 suite(37). The ff12SB and Glycam-06 force fields were used for the protein and oligosaccharides, respectively(37). The simulation protocol used corresponds to the one described in our previous paper(35). It basically consisted in a series of minimizations and MD equilibration steps in the NPT and NVT ensembles to obtain a series of equilibrated initial conformations. Each of those conformations was subjected to eight independent NVT simulations lasting 50 ns each.

#### 2.2.3 Hydrodynamic calculations

We used the BEST procedure discussed in the Introduction. A total of 25 atomistic structures picked up from each of eight independent trajectories for each simulated system were submitted to BEST. This procedure mainly consists of three steps(38). In the first step one uses the procedure described by Connolly where one obtains the accessible surface to solvent by rolling a probe particle around the protein structure(39). In our case we used a 1.5 Å probe radius to mimic the water molecule. In a second step, a surface triangulation is carried out and refined by a coalescent post-processing method to get data suitable for boundary element calculation. The last step is the calculation of hydrodynamic properties by solving the large linear system of equations with the stick boundary conditions. This step is the most computer-demanding part of the calculations as it scales as the cube of the triangle number used in the triangulation algorithm. The strategy suggested by Aragon was followed in this case. It consists of performing calculations with different number of triangles and extrapolating to an infinite number of them. Averaged quantities for the translational diffusion coefficient, D, and the intrinsic viscosity [η] were reported for each of those 200 protein conformations (25 structures from each of the eight independent MD runs). All calculated values are referred to T = 293 K with a solvent viscosity of 1.002 mPa·s. Details of the methodology employed are given in the Supplementary Information section of ref. (35).

## 3. Results

### 3.1 Molecular sizes and hydrodynamic properties

#### 3.1.1 SEC experiments

Absolute molecular weights (M_w_), intrinsic viscosity ([η]) and hydrodynamic radius (r_h_) have been obtained from the SEC combining the tetradetection experiments. Figure 2 shows chromatograms obtained for the g-eHER2-his, TZM solutions as well as the 2:1 and 1:2 TZM:g-eHER2-his mixtures. Absolute molecular masses of 88.7 ± 1.8 kDa and 149.0 ± 3.0 kDa were measured for g-eHER2-his and TZM, respectively. The measured molecular mass of g-eHER2-his indicates mainly a monomer form in solution under the experimental conditions used. On the other hand, the observed TZM molecular mass value is in agreement with the typical size for antibodies of the IgG type (40).

The appearance of peaks at shorter retention times for both 2:1 and 1:2 TZM: g-eHER2-his mixtures with respect to the control samples (Figure 2) are indicative of the formation of complexes. In the 2:1 mixture with an excess of TZM, a peak at 11.7 ml is resolved, giving an absolute molecular weight of M_w_ = 245.0 ± 5.0 kDa. This molecular weight is consistent with a hetero-dimer of one TZM with one g-eHER2-his (1:1 complex). The peak corresponding to the unbounded TZM (at 13.6 ml) is also observed. Moreover, a shoulder appears at lower retention volumes, around 10.7 ml, giving rise to a molecular weight of M_w_ = 349 ± 13.0 kDa, that is consistent with a hetero-trimer of one TZM and two g-eHER2 (1:2 complex). This peak is well resolved in the 1:2 mixtures with an excess of g-eHER2, as it can be observed in Figure 2. The peak of the unbound g-eHER2-his (at 13.2 ml) is found as well.

The intrinsic viscosity [η] can be calculated using the well-known viscosity Einstein relationship:

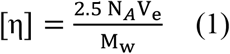

where M_w_ is the molecular weight, N_A_ is the Avogadro’s number, and V_e_ is the volume of an equivalent spherical particle. In this equation the hydration shell around the molecules are defined in terms of equivalent hydrodynamic spheres that would increase its viscosity to the same extent as a solid spherical particle of volume V_e_=4/3πr_h_^3^, resulting in the following equation for the hydrodynamics radius:

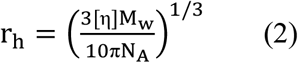

**Fig. 2.**
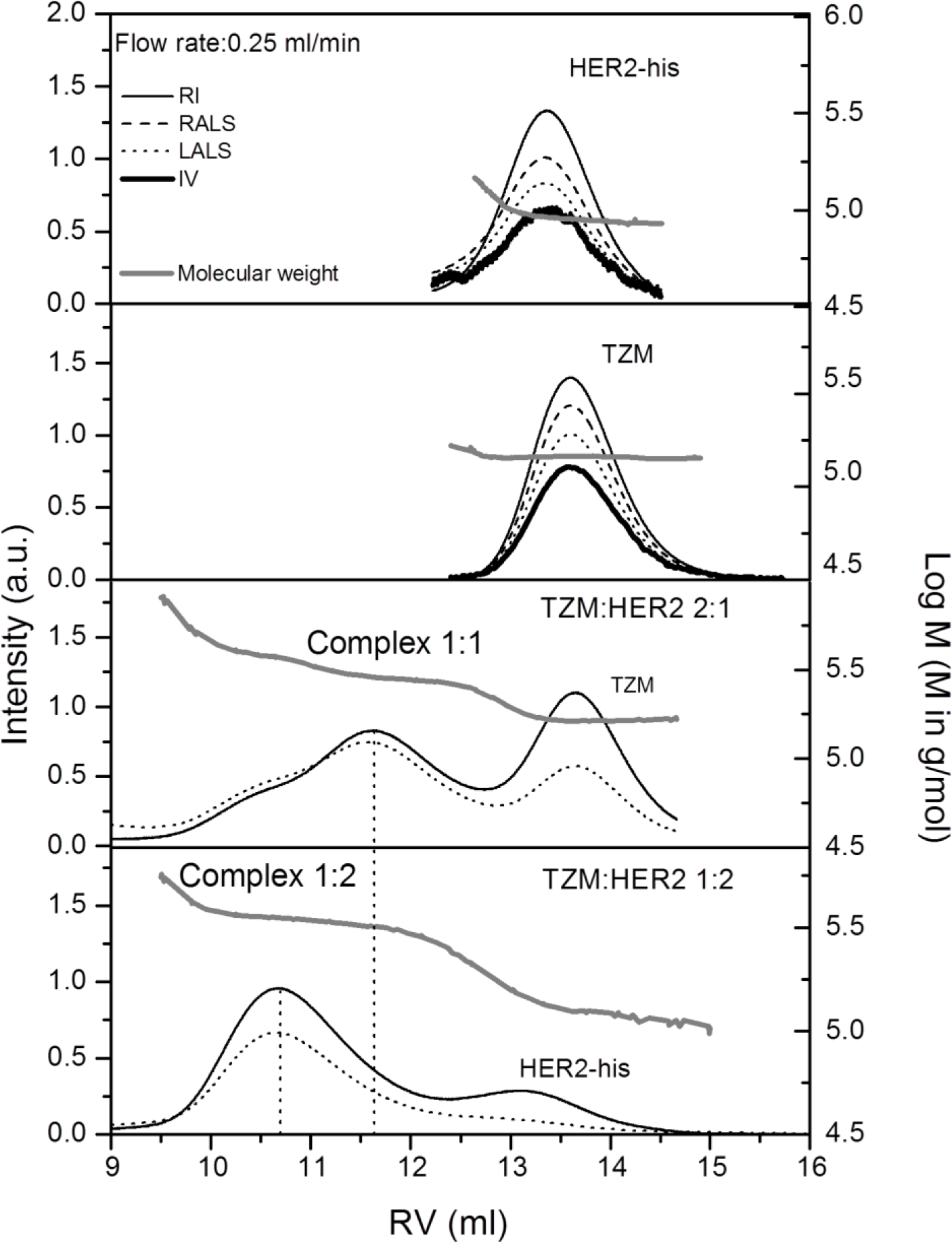
SEC/tetradetection chromatograms of TZM, g-eHER2-his, complex 1:1 (hetero-dimer) and complex 1:2 (hetero-trimer).

Both [η] and r_h_ values are listed in Table 1. The [η] for the bounded complex 1:1 and 1:2 are 7.4 and 8.6 cm^3^·g^−1^, respectively. These values are higher than those observed for globular proteins. In general non-glycosylated globular proteins show values of [η] located between 2.5 and 4.5 cm^3^·g^−1^ (see Figure 7 in reference(35)). Furthermore, they are even larger values that those found for the unbounded species (TZM and g-eHER2-his, …). This is likely due to two factors: i) a greater flexibility of the protein and ii) a larger number of glycans in the bound complexes. Both factors have been reported to play a role in the protein hydrodynamics.(35, 41, 42)

In fact, our previous MD simulations performed on the members of the EGFR family in water, detected a large root mean square fluctuation concentrated in domains I, II and IV of the systems, indicating the existence of different movements of the domains and hence a high degree of flexibility (43–45). Also, using MD simulations, we observed a profound effect of glycan chains on hydrodynamics properties of the e-HER2 protein. (35)

**Table 1.** E*xperimental molecular and hydrodynamic results for the structures studied at*

| Sample | M<sub>w</sub> (kDa) | [$\eta$] (cm<sup>3</sup>·g<sup>-1</sup>) | r<sub>h</sub> (nm) | D×10<sup>7</sup> (cm<sup>2</sup>·s<sup>-1</sup>) | r<sub>h</sub> (nm) |
| --- | --- | --- | --- | --- | --- |
|  | SEC | SEC | SEC<sup>a)</sup> | DLS | DLS |
| *g-eHER2-his*<sup>b</sup> | 88.7 ± 1.8 | 6.4 ± 0.2 | 4.5 ± 0.1 | 4.46 ± 0.02 | 4.7 ± 0.1 |
| *TZM*<sup>b</sup> | 149.0 ± 3.0 | 6.5 ± 0.1 | 5.4 ± 0.1 | 4.01 ± 0.02 | 5.2 ± 0.1 |
| *Complex 1:1* | 245.0 ± 5.0 | 7.4 ± 0.3 | 6.8 ± 0.1 | 3.16 ± 0.02 | 6.6 ± 0.1 |
| *Complex 1:2* | 349.0 ± 13.0 | 8.6 ± 0.3 | 7.7 ± 0.1 | 2.50 ± 0.02 | 8.4 ± 0.1 |

<sup>a</sup> The hydrodynamics radius has been calculated using the equation 2 with the data collected from the chromatograms, <sup>b</sup> data from Vega et al. 2017, BBA-General Subjects.(35)

#### 3.1.2 DLS experiments

Additionally, [η] and D can be obtained by means of DLS experiments. The r_h_ values can be calculated from DLS taking the D values by using the Stokes-Einstein relationship. DLS results are presented in Fig. 3 as the squared electric field autocorrelation function, [g_1_(t)]^2^, for the complexes at T = 293 K. Results obtained previously for TZM control are included for comparison purposes.(35) A clear delay of [g_1_(t)]^2^ is observed as one moves from TZM to complex 1:1 and 1:2, in agreement with an increase of the molecular size of the complexes.

The cumulate analysis described also in reference (35) has been proved to be suitable for obtaining the mean translational diffusivities of the complexes from the DLS autocorrelation functions, and the results are listed in Table 1.

**Fig. 3.**
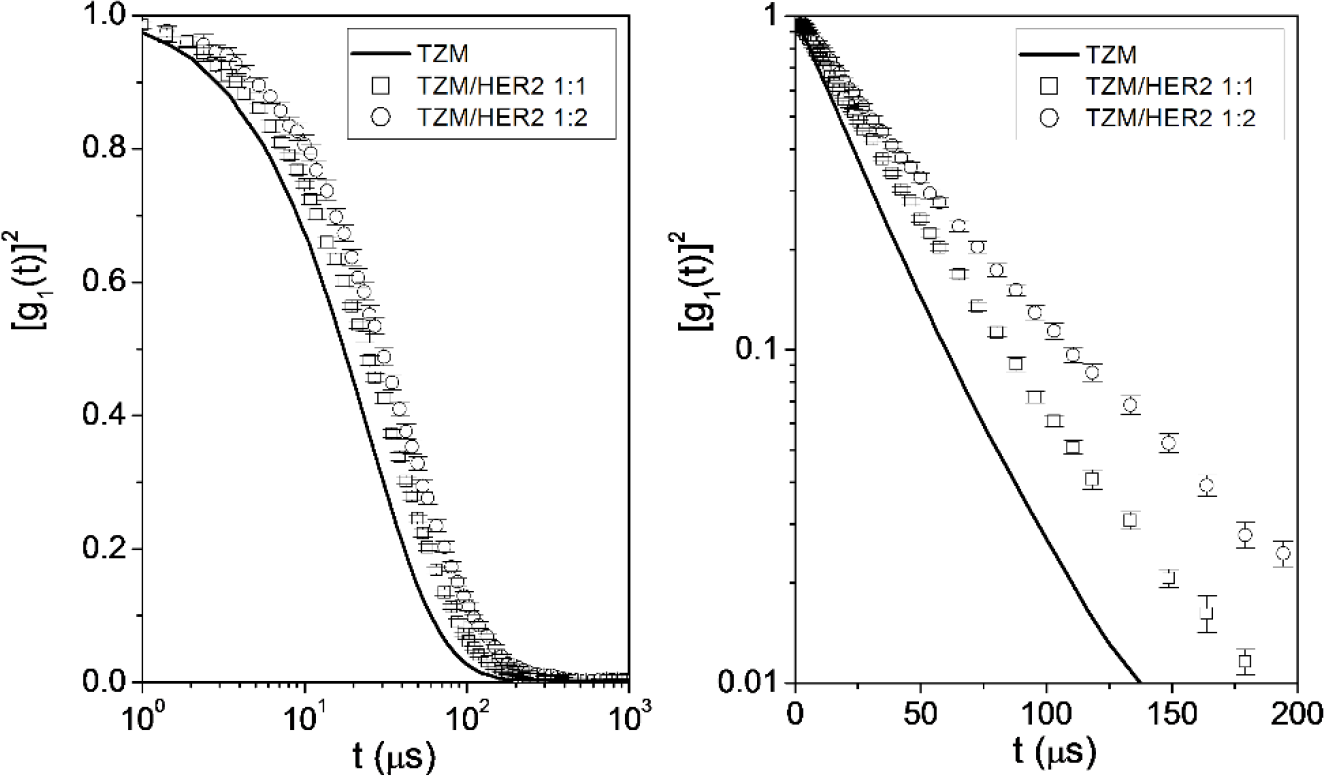
Squared electric field time autocorrelation function, [g_1_(t)]^2^, of TZM (solid line), complex 1:1(◻), and complex 1:2 (◯) at T = 293 K

We have plotted in Fig. 4 the results for r_h_ obtained from DLS experiments as a function of the experimental molecular sizes. In the figure the results are compared with the reported behavior observed in globular proteins (34, 46, 47). It can be clearly observed the deviation obtained for g-eHER2-his, TZM and their complexes, which can be explained by the lower molecular density, [η]^−1^, most probably induced by the glycans and the intrinsic flexibility of both g-eHER2 and TZM, as explained in the previous section.

**Fig. 4.**
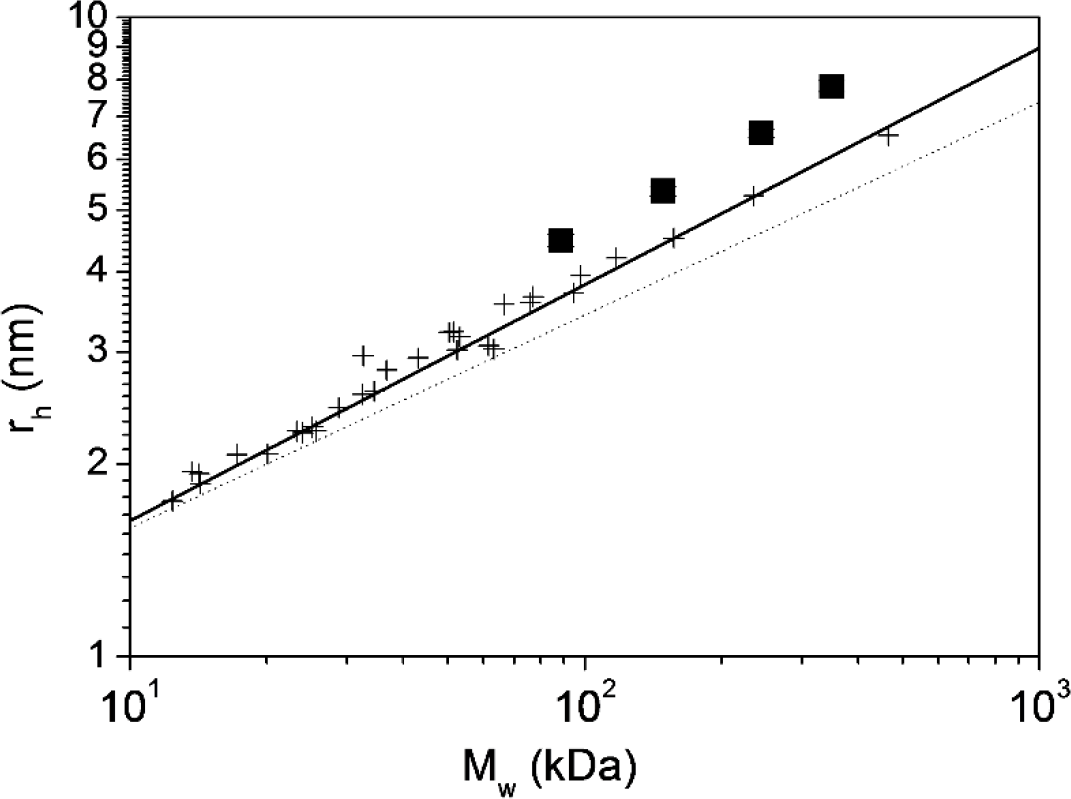
Hydrodynamic radius, r_h_, versus molecular size, M_w_, for the systems under study (■) and globular proteins (+) from the literature (34, 46, 47)

#### 3.1.3 Electrostatic properties

Charged groups can certainly play an important role in biochemistry and biophysics of proteins. Electrostatic interactions have a strong influence in processes as selective transport in protein channels, folding and denaturation, crystallization, association of receptors with ligands, or, as in the present study, formation of protein-protein complexes (48). Thus, the determination of electrostatic properties in our systems is in principle a very valuable tool to characterize the interactions between g-eHER2-his and TZM.

The results of the electrophoretic mobility, μ_e_, obtained in the EM measurements for the samples under study are given in Table 2. The EM measured can be used to determine the ζ-potential using Henry’s equation:

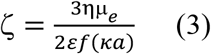

where ε is the dielectric constant of the medium, η is the viscosity of the dispersant, ζ is the zeta potential and f(κa) is the Henry’s function. The values of f(κa) have been estimated using the approximate expression given by Swan and Furst (49). If the values of ζ of a particle is less than k_B_T/e (i.e., 25.7 mV at 298 K), the effective molecular charge can be evaluated using the Debye-Hückel-Henry (DHH) approximation to correct for ionic radii and ionic strength effects:

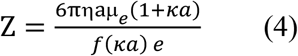

where e is the electronic charge, a is the particle radius, and κ the inverse Debye length. This later can be calculated for monovalent salt at T = 298 K at any ionic strength, I, as:

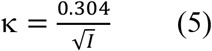

The radius of the particle has been taken as the hydrodynamic radius, r_h_, obtained from DLS measurements in Table 1 and from reference for g-eHER2 (35).

**Table 2.** Electrostatic properties of the systems under study

| *Sample* | $\mu_e$<br>($\mu\text{m}\cdot\text{cm}\cdot\text{V}^{-1}\cdot\text{s}^{-1}$) | $\zeta$<br>(mV) | $Z$<br>(EM) | $Z$<br>(PropKa/Delphi)<sup>a</sup> | $\Delta Z^b$ | pI |
| --- | --- | --- | --- | --- | --- | --- |
| *g-eHER2* | $-0.76 \pm 0.04$ | $-9.1 \pm 0.5$ | $-18.3 \pm 1.4$ | $-17.0 / -16.6$ | 1.3 / 1.7 | 5.9/6.3 |
| *TZM* | $+0.28 \pm 0.02$ | $+3.3 \pm 0.3$ | $+8.3 \pm 0.9$ | $+10.0 / +12.9$ | 1.7 / 4.6 | 8.8/9.7 |
| *Complex 1:2* | $-0.45 \pm 0.03$ | $-5.0 \pm 0.3$ | $-26.1 \pm 1.7$ | $-22.0 / -19.9$ | 4.1 / 6.4 | 6.7/6.8 |
| *Complex 1:1* | $-0.29 \pm 0.01$ | $-3.3 \pm 0.1$ | $-13.2 \pm 0.6$ | $-4.0 / -3.1$ | 9.2 / 10.1 | 7.3/7.2 |

<sup>a</sup> PROPKA3 within the PDB2PQR server [43,44] and DelphiKa [45] packages were used to calculate these values. <sup>b</sup> $\Delta Z$ is the difference between the experimental charge and those calculated by simulations

The estimated values ζ-potential and Z using Eqs. (3) - (5) are listed in Table 2 for the systems under study. In general it is considered that the values obtained for the effective charge, Z, will be close to the “true net charge” of the proteins and complexes (48). Notwithstanding, it should be emphasized that within the framework given by Eqs (3) – (5), Z is the fixed charge arising not only from the ionized groups on the protein, but also from the ions bound in the Stern layer.

### 3.2. MD Simulations and Hydrodynamic Modeling

#### 3.2.1 TZM:g-eHER2-his interactions

Figure 5 shows the residue-residue contact maps between amino acids corresponding to the monoclonal antibody Fab and g-eHER2-his domain IV either in 1:1 or 1:2 complexes obtained from the MD simulations. The experimental crystallographic structure is also shown (PDB ID: 1N8Z). This map has been calculated using the MDcons tool(50). Each point means a residue contact with a persistent distance (larger than 90% in the MD trajectory) below 5 Å between the corresponding centers of mass. The interactions of the crystallographic structure are largely preserved along the molecular dynamics simulations. A few differences can be observed with respect to the contacts present in the crystallographic structure. For instance, some interactions between the loop containing residues 570 to 573 in domain IV with the Fab heavy chain and contacts between residues 598-600 and the light chain are lost in the simulated systems. In principle, this fact cannot be attributed to the presence of glycans on residues ASN549 or ASN607 due to the outwards orientation of the sugar groups. It may be due to the presence of the histidine tag at the C-terminus in the simulated system. In spite of these slight differences, it can be concluded that the residue-residue contacts are quite similar in simulated solution models and in the crystallographic structure.

**Fig. 5.**
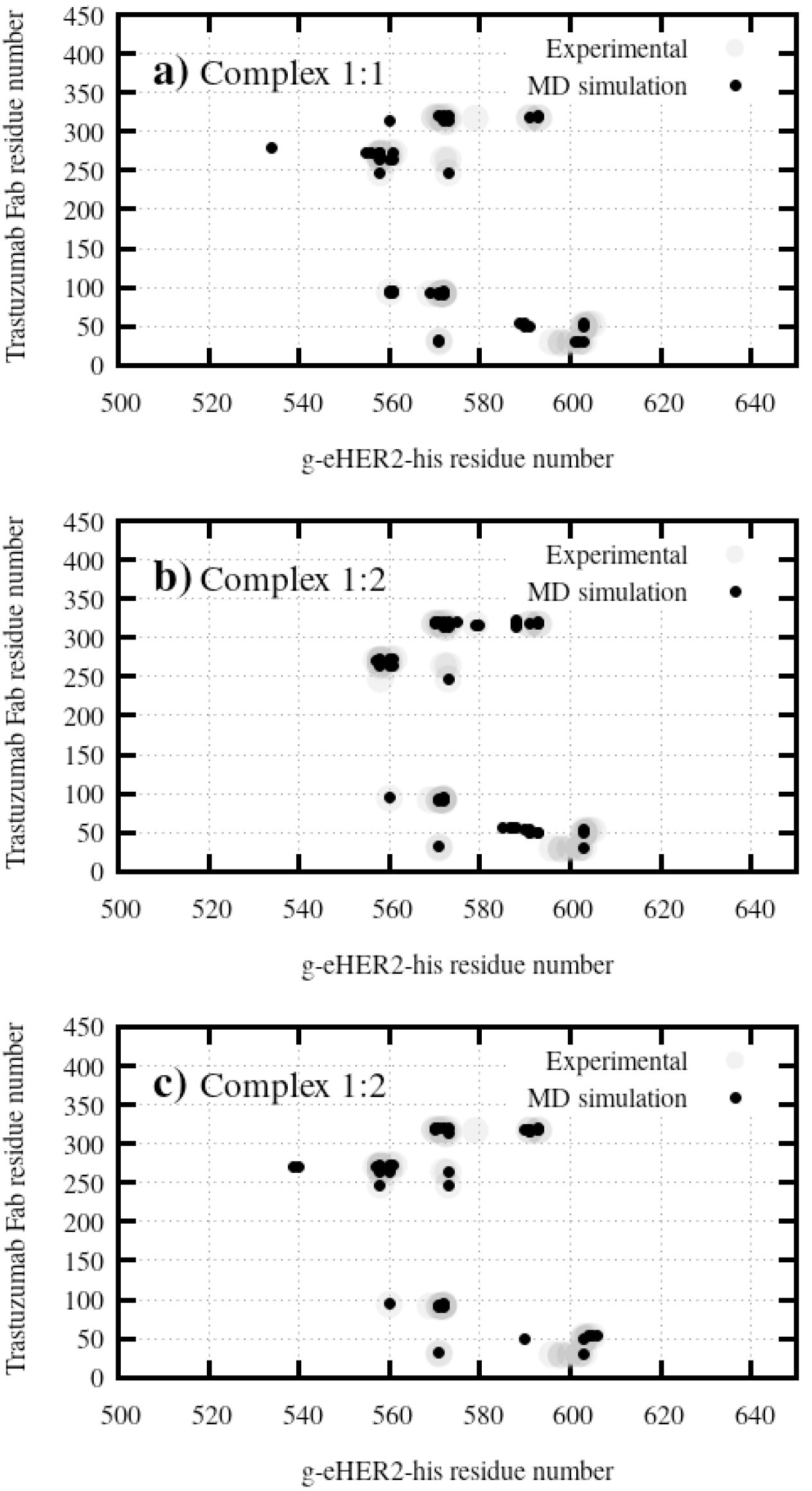
Residue contact maps corresponding to: **a)** the simulated complex 1:1 and **b)** an **c)** correspond to the interaction between each g-eHER2-his molecule in the 1:2 complex and the corresponding Fabs in the TMZ compared to the crystallographic data (PDB code:1N8Z). The contact maps were calculated using the MDcons software(50). Only contacts with a presence larger than 90% in the MD trajectory are shown.

Figure 6 shows a representative snapshot extracted from the NVT simulations and used to calculate the hydrodynamic properties. It can be observed the ability of the monoclonal antibody to bind two g-eHER2-his extracellular receptors without perturbing the interaction between any of them with its corresponding Fab fragment. The relative mutual orientation of the two Fabs and the Fab-domain IV interaction geometry provides enough space to avoid steric congestion between the two g-eHER2-his structures.

**Fig. 6.**
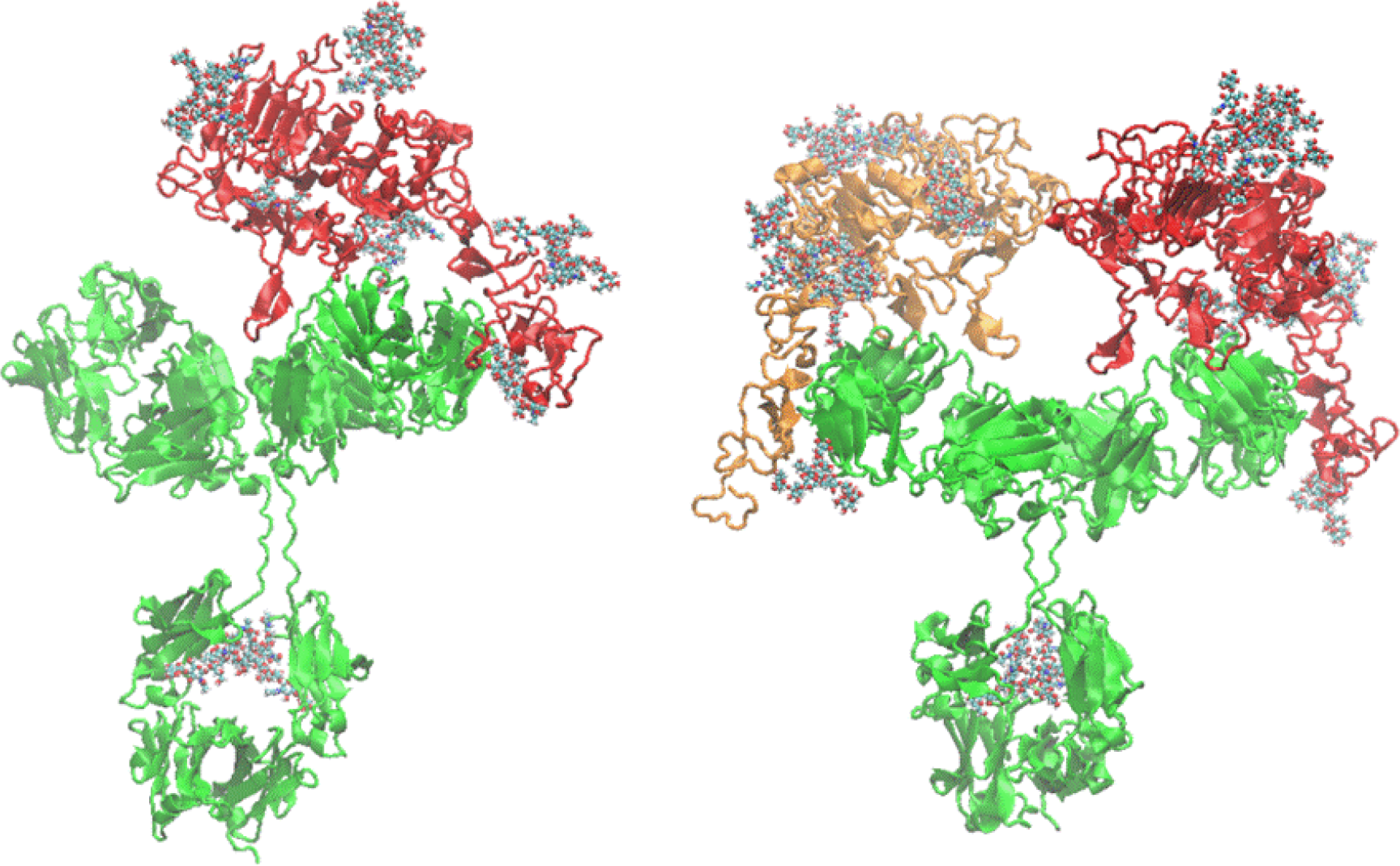
Cartoon representation of representative equilibrated conformations for the 1:1 (left) and 1:2 (right) complexes. The TZM antibody protein is represented in green and the g-eHER2-his in red and orange colors. The glycans are represented as CPK models. PDB files of these structures can be found in the supporting material.

#### 3.2.2 Hydrodynamic properties calculation

The translational diffusion coefficient, hydrodynamic radius and intrinsic viscosity were calculated following the procedure proposed by Aragon (38). The number of triangles considered in the coalesce datastep varied from 6000 to 15000 for the 1:1 complex and from 8000 to 17000 for the 1:2 complex. These values represent a compromise between accuracy for the extrapolation of the hydrodynamic values and the computational efficiency. The extrapolation to infinite number of triangles was supported by a linear fit of the dependent variable against 1/N, being N the number of triangles. Correlation coefficients larger than 0.9 are obtained for those linear fits. The total molecular weights calculated for complexes 1:1 and 1:2 are 234.6 and 320.3 kDa, respectively. These values were used to evaluate the hydrodynamic quantities in each case. The averaged values obtained for the intrinsic viscosity [η] were 7.9 ± 0.2 (cm^3^·g^−1^) and 8.4 ± 0.1 (cm^3^·g^−1^) for complexes 1:1 and 1:2, respectively. The translational diffusion coefficients (D) averaged over all the trajectories were 3.18 ± 0.02 ×10^7^ (cm^2^·s^−1^) and 2.80 ± 0.01 × 10^7^ (cm^2^·s^−1^) for the 1:1 and 1:2 complexes, respectively. The computed results will be compared in the next section to those obtained experimentally.

## 4. Discussion

The comparison between the calculated and experimental hydrodynamic properties can be observed in figure 7. To enrich the discussion, data taken from the literature (mostly globular monomeric and multimeric proteins) will be also included. As it can be observed there is a nice agreement between the computational hydrodynamic analysis obtained from the MD trajectories and the experimental measurements for the systems under study. The percentage difference between the calculated and experimental values are less than 5 %, with the exception of the diffusion coefficient of the complex 1:2, which is almost 10%. These results support the fact that the used approach works very well, not only in the cases of flexible antibodies and globular proteins (34, 46, 47), but also for heavily glycosylated flexible proteins such as monomeric, homodimeric and antibody-antigen complexes of different nature, as studied in the present work.

As it can be observed, the values of [η] (Fig. 7a), in our case solid symbols, fit very well with the experimental values, and they are nearly twice as those corresponding to globular proteins. The values of D obtained are the lowest ones within the range explored in previous works, and significantly lower than those expected for globular systems with the same molecular weight (Fig. 7b).

Concerning electrostatic properties in Table 2, the reported positive value of Z for TZM at pH 7.5 is in the same range as those values reported in other studies for IgG1 antibody at pH within 5.0 and 9.0 and low ionic strength (23, 51–54). The result obtained for TZM are in contrast with the high negative charge density of the g-eHER2 surface, as judged by the results shown in Table 2, presumably due to exposed aspartic and glutamic acids residues. Changes in either the sign or the magnitude of the ζ-potential of these systems can serve as a qualitative indication of changes in the surface charge density, e.g., specific interactions between the biomacromolecules (55). Interestingly, the complexes 1:1 and 1:2 show intermediate values of the ζ-potential. These intermediate values indicate a neutralization of the charge at the surface of the complexes, also confirming that they consist of both TZM and g-eHER2-his proteins.

**Fig. 7.**
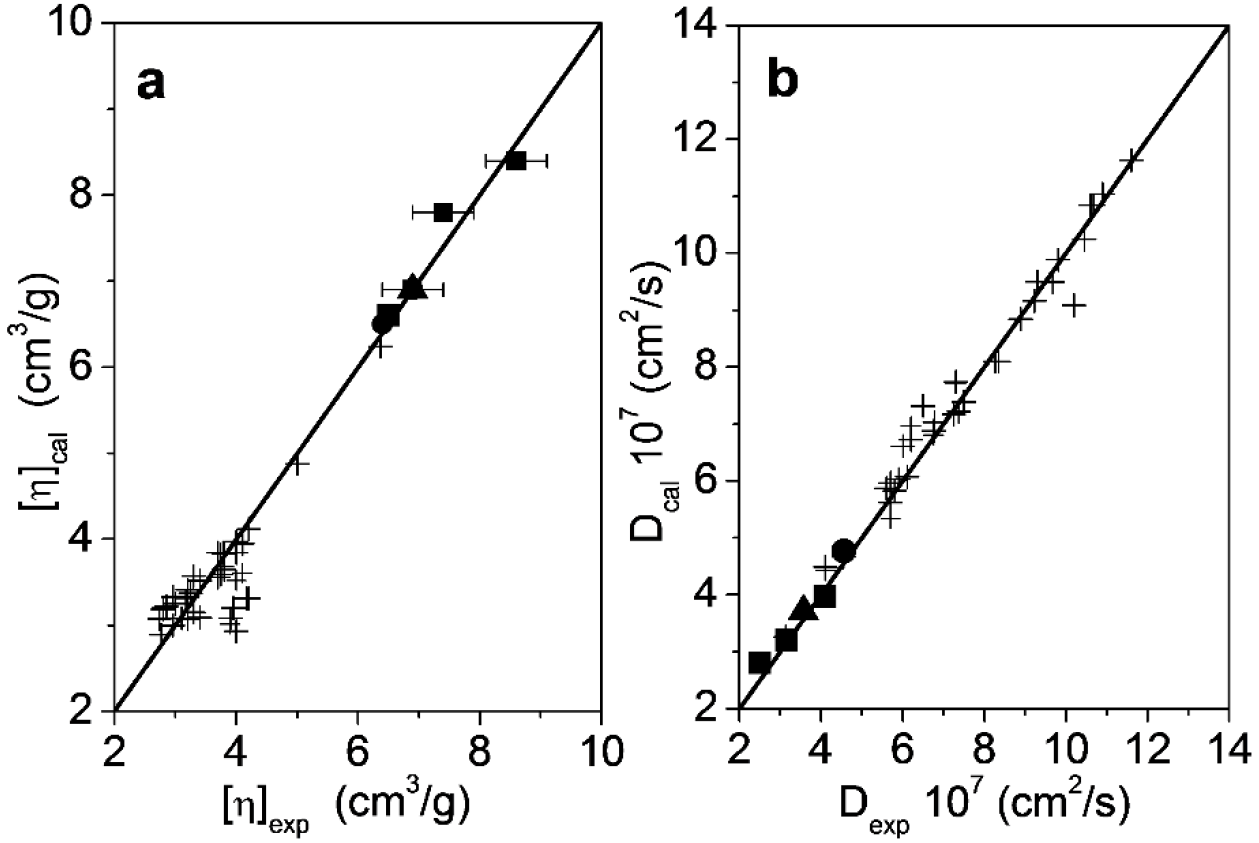
Comparison between calculated and experimental values of intrinsic viscosity and translational diffusion coefficient of the systems under study, (■) TZM and complexes, (●)g-eHER2 monomer [19] (▲) g-eHER2 dimer [19], and (+) mono and multimeric globular proteins taken from the literature (34, 46, 47)

The values for the net charge for the systems under study have also been calculated theoretically using PROPKA3 within the PDB2PQR server (56, 57) and Delphi pKa (58) packages. The agreement between experiment and theoretical calculations is especially good in g-eHER2-his sample (ΔZ = 1.3 − 1.7). In the case of TZM slightly higher values for theoretical charge is obtained when compared to the experiments (ΔZ = 1.7 − 4.4). Theoretical values differ significantly from the experiments in the 1:2 complex (ΔZ = 4.1 − 6.4), but especially in the 1:1 complex (ΔZ = 9.2 − 10.1). Nevertheless, the net charge ranking in experiments and theoretical calculations is consistent, as it can be observed in Fig. 8.

It has been reported that below or in the vicinity of the isoelectric point (pI), where a protein carries a positive charge, the measured value of Z is lower than that obtained from theoretical calculations due to the Hofmeister effect or the specific binding of anions to the proteins (59–62), even at low ionic strength (63, 64). For example, in NaCl solutions of BSA, the excess number of bound chloride ions ranged from 40 at pH 3 to 5 ions at pH 6.5, this later above of the isoelectric point of BSA, pI 5.3. In fact some bound chloride is found at pH as high as 8.0 in this case (65).

**Fig. 8.**
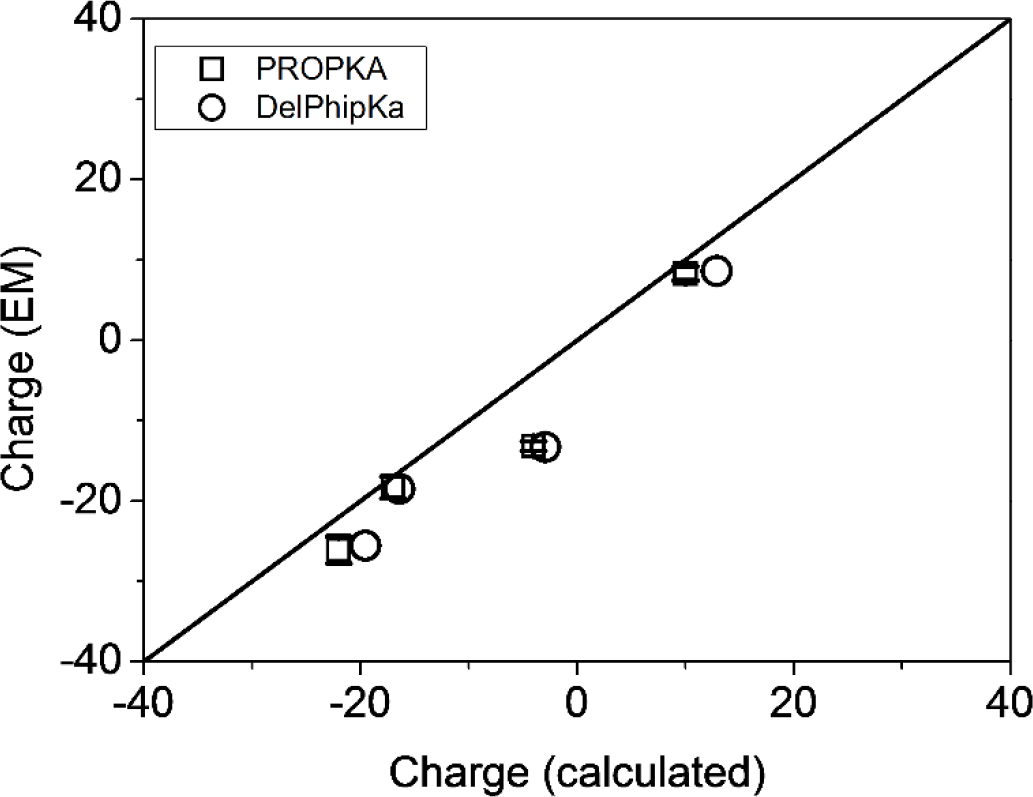
Net charge for the systems under study (experimental *vs* theoretical data)

It should be noted that the calculated values from PROPKA and Delphi pKa packages do not account for the charges arising from ions bound in the Stern layer, which can substantially contribute to the measured ζ-potential and Z. In the case of g-eHER2-his sample the difference between experimental and theoretical charge values (ΔZ = 1.3 − 1.7) is quite close to the experimental uncertainty of the ζ-potential and Z, or may be also due to the use of different force fields for the theoretical calculations. In addition, the experimental pH 7.5 is nearly 2 units above the estimated pI 5.8 for this protein from the calculations, so no chloride binding is expected. However, some degree of preferential anion binding may occur in the TZM (ΔZ = 1.7 − 4.6) at pH 7.5, well below the TZM pI 9.1, recently measured experimentally (66). This result is also in agreement with fresh experiments that revealed specific anion effects at low ionic strength in mAbs solubility behavior (23, 51, 52). Thus, the slightly lower experimental Z value relative to the theoretical charge in TZM sample may indicate a preferential binding of chloride anion to the positively charged protein groups.

In the case of 1:2 complex the difference between experimental and theoretical charge values is also modest (ΔZ = 4.1 − 6.4). Thus the difference may still be explained due to a similar amount of chloride binding than in TZM, as the theoretical pI 6.8 in 1:2 is close to the experimental pH 7.5. A similar deviation would be expected for the 1:1 complex, as, like 1:2, it is composed also by one TZM unit. However, in this case the deviation from the theoretical value is twice (ΔZ = 9.2 – 10.1) the value that is observed for TZM or 1:2 samples.

**Fig. 9.**
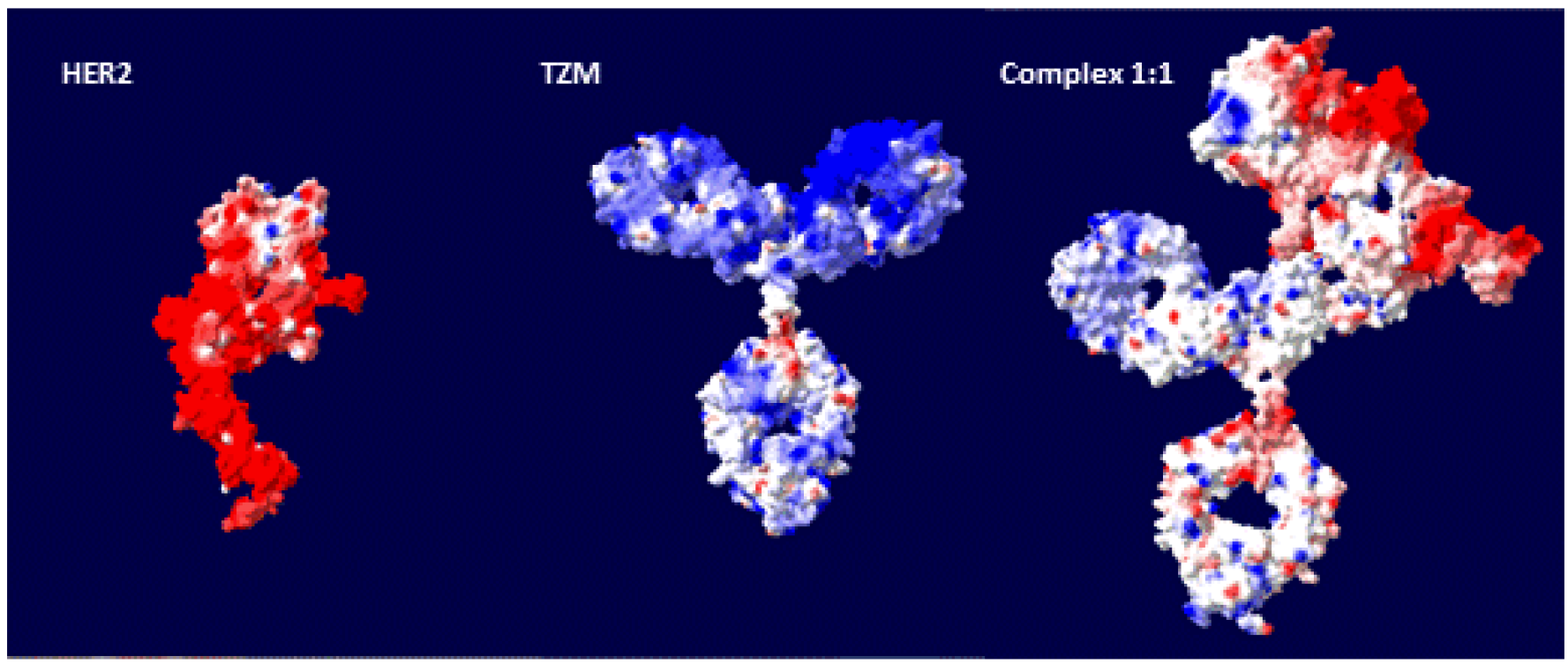
Distribution of the electric charge at the molecular surface of the systems under study. The molecular surface is colored as follows: red (negative cutoff −2.0 k_B_T/e), white (neutral points), blue (positive cutoff +2.0 k_B_T/e) color gradient. The simplest calculation for the electrostatic potential (Coulomb) has been selected as a first approach, using the Deep View (Swiss-PdbViewer) package (67)

For a better visualization of the electrostatic nature of the surface of the systems, we have included in Fig. 9 a calculation of the charge distribution for the g-eHER2-his, TZM and complex 1:1 using the DeepView (Swiss-Pdb Viewer) package. (67) It is clearly seen the highly asymmetric charge distribution in the complex 1:1. It is worthwhile to mention here that Debye-Hückel-Henry (DHH) approach [Eq. (4)], is based on the uniform surface charge density model, that is in fact a general formula for a sphere with an arbitrary charge distribution. However, this approach is widely applied to EM data in proteins, since a uniform surface charge density model is hardly applicable. There are theoretical approaches that provide explanations about possible changes in electrostatic features that go beyond a simple counting of the number of charged residues on the surface of the particles. For spherical and ellipsoidal Brownian particles with a non-uniform charge distribution a simple interpretation in terms of the total charge is a matter of debate, mainly because the μ_e_ value may depend on the magnitudes and special orientation of multipole moments of all orders (mainly total charge, dipoles and quadrupoles), even in ellipsoidal and spherical particles (68–70). Actually, the theoretical approaches predict measurable μ_e_ in the case of neutral particles with a non-uniform surface charge distribution.

In addition, it should be noted that unlike rigid charged colloids, for which a straightforward relation between electrostatic properties and surface charge exists, proteins could be considered as “soft particles” (71, 72). In these systems electrostatic properties are dependent not only on the net charge on the surface, but also on charge distribution and permeability with respect to solvent and ions. The value of μ_e_ is determined, in this case, by a subtle balance between the electrical force and electro-osmotic drag. In particular, the theoretical model developed by Duval et al. for diffuse soft particles shows that the screening of the electro-osmotic drag increases upon increasing the charge density and hydrodynamic permeability, thus resulting in a higher magnitude of μ_e_.

In a different framework, by considering again the case of hard spheres and the DHH approximation, Lošdorfer-Božič and Podgornik (73) analyze how the symmetry of charge distribution modifies the interaction energy. They found that local charge inhomogeneities affect on a different way the electrostatic interactions of equally charged particles. More recently these authors have established a methodology for extracting the different multipole moments (net charge, dipoles and quadrupoles) accounting only for the exposed residues in globular proteins. They show that not only these magnitudes, but more importantly spatial orientation, have profound effects on electrostatic interactions (74). A closer inspection about these features in the case of the complexes studied here would be desirable, as the highly asymmetric charge distribution will have consequences for the study of protein-protein interactions.

## 5. Conclusions

In this work, we have performed a combination of experiments, modeling and simulation techniques to investigate dynamics, transport and electrostatic properties of the complexes formed by trastuzumab and glycosylated HER2 extracellular domain in aqueous solution. These complexes are relevant in the treatment of HER2++ cancer with monoclonal antibodies. There is a very good agreement between experimental and calculated hydrodynamics properties. The higher values of the intrinsic viscosity and lower values of the diffusion coefficient, with respect to those corresponding to globular proteins of similar molecular mass, supports a high degree of molecular flexibility in both the dimer (complex 1:1) and the trimer (complex 1:2)

The simulations indicate that the complexes in solution are firmly bound, as indicated by similarity of the residue-residue contacts in the interaction surface between the TZM and the g-eHER2-his proteins in the simulated solution models and in the experimental crystallographic structure. However, they retain, in other parts, the interesting and specific dynamic features of the monomeric TZM and g-eHER2 species. This high flexibility is caused, on one hand, by the open conformation of the receptor and on the other hand, by the large root mean square fluctuations of the different domains, especially the g-eHER2-his domain IV. This is probably due to its hinge movement, previously reported by us (35, 43, 44). In addition, a highly asymmetric charge distribution is detected for the 1:1 TZM/HER2 complex, which has strong implications in the correlations between the experimental electrophoretic mobility and the modeled net charge using computational tools.

## Supporting Material

A zip file containing the PDB structures shown in Figure 6 is available.

## Authors contributions

J.M.S. and J.C. designed the research; J.R. and V.C. performed the computer simulation research; J.F.V. and E.S.-S. performed the experimental research, J.C. provided new reagents; J.M.S., J.R., J.F.V., and V.C. analyzed the data; and J.M.S., J.R., J.F.V., and V.C. wrote the article.

## Acknowledgments

This research work was supported by the Spanish Ministry of Economy and Competitiveness (MINECO, Spain) (Project MAT2012-36341-FEDER) and by the CSIC (Spain) - Project PIE201360E097. J. Ramos acknowledges financial support through the Ramón y Cajal Program (MINECO, Spain) - Contract RYC-2011-09585. We acknowledge Prof. Sergio R. Aragón for kindly providing us the BEST software to compute the transport properties of biomacromolecules. There are no conflicts to declare

